# Epigenetic transmission of cardiometabolic risk in offspring discordant for maternal gestational metabolic fitness

**DOI:** 10.1101/435461

**Authors:** Frédéric Guénard, Simon Marceau, Serge Simard, John G Kral, Marie-Claude Vohl, Picard Marceau

**Author notes:** These authors contributed equally to this work (John G Kral, Marie-Claude Vohl, Picard Marceau). **Corresponding Author:** Marie-Claude Vohl, Ph.D., Institute of Nutrition and Functional Foods (INAF), Laval University, 2440 Hochelaga Blvd, Quebec, Quebec, Canada, G1V 0A6.

## Abstract

**OBJECTIVE:** Diabesity during gestation predisposes offspring to lifetime cardiometabolic risk. We demonstrated that maternal biliopancreatic diversion surgery (BPD) improved metabolic fitness and pregnancies reducing cardiometabolic risk in siblings born after (AMS) compared to before maternal surgery (BMS). We found differential methylation of inflammatory and glucoregulatory genes in peripheral blood cell DNA in offspring and in mothers after BPD compared to preoperative “control” women. Here we study offspring trajectories of cardiometabolic risk markers and determine persistence of the methylome related to body weight (BMI) and gestational weight gain (GWG).

**RESEARCH DESIGN AND METHODS:** Prospective, cross-sectional study of 133 mothers with 89 BMS and 183 AMS offspring born mean 4 years after BPD and 83 unoperated control women was conducted, and differential methylation patterns in mothers were compared with those of offspring during 2-26 years.

**RESULTS:** Independent of maternal or offspring BMI and GWG, postoperative maternal metabolic fitness was associated with improved cardiometabolic phenotype in AMS vs. BMS offspring sustained beyond puberty. BMS offspring exhibited increasing linear trajectories of weight, cardiometabolic and inflammation risk factors *versus* normative horizontal trajectories of AMS offspring. Methylation differences between AMS and BMS offspring identified 45 625 differentially methylated sites, 73% overlapping with those of mothers vs. controls; 4 446 demonstrated similar sustained directionality of differences in methylation levels; 154 sites exhibited significant correlation coefficients (r≥0.4) overrepresented within genes associated with cardiometabolic risk, growth and inflammation.

**CONCLUSION:** Maternal BPD appears to epigenetically prevent transmission of cardiometabolic risk independent of BMI.

---

Diabesity, a chronic inflammatory insulin-resistant dysmetabolic syndrome is increasing world-wide, with no evidence of effective prevention (1). Trans- or inter-generational transmission of diabesity is well documented in pre-clinical and clinical investigations identifying environmental and genetic contributions of both parents. Studies of mechanisms have mostly focused on environmental factors with very few on the preventive or therapeutic efficacy of intervention in humans, owing to reluctance to manipulate energy balance during gestation related to historic and evolutionary memory of adverse outcomes during times of famine (2,3). Restricting maternal intake (diet) and/or increasing exertion (exercise) in metabolically unfit women to limit excessive gestational weight gain (GWG), with its well-known hazards (4), have largely been ineffective; careful drug studies are controversial.

The increasing practice of surgical treatment of diabesity the last half-century has created an observational fund of knowledge of pregnancy outcomes after maternal operations (5). These operations, although collectively termed “bariatric”, vary, not only in their mechanisms, but also in the durability of their effects. In distinction to purely restrictive banding, most effective currently practiced laparoscopic operations are hybrid such as gastric bypass (RYGB), and pylorus-preserving biliopancreatic diversion (BPD) (6) and the neuroendocrine, restrictive sleeve gastrectomy. Within-subject comparisons have enabled relatively well-controlled studies exhibiting the improved risk-benefit ratio of undergoing preconception “bariatric” surgery especially with respect to maternal gestational health, complications, fetal loss, delivery and neonatal health although there have been lingering concerns over the potential adverse outcomes of malabsorptive operations (7). Less studied are long-term outcomes of offspring born after maternal surgery.

Numerous surveys have documented relationships between birth weight, neonatal weight gain and trajectories of weight development during different “critical periods” and adult cardiometabolic diseases (8,9). Growth trajectories of childhood obesity exhibit sharp increases in prevalence carried into adulthood (10) associated with high risk of type 2 diabetes at ≥30 years in men (11). Reflecting the durable beneficial effects of metabolic surgery in reproductive-age women, we described similar benefits in offspring born after (AMS) compared to those born before (BMS) biliopancreatic diversion (12,13), using the prospective Université Laval Mother-Child Obesity Study (M-COS). Postulating that these differences were related to maternal metabolic fitness affecting the germline and/or intra-uterine environment, we studied epigenetic differences in blood DNA comparing BMS to AMS offspring (14,15) and their mothers to matched preoperative women (16).

Here we present differences and correlations between trajectories of cardiometabolic risk markers out to 26 years of age and differential methylation (epigenetic markers) of cardiometabolic genes in offspring discordant for maternal metabolic surgery.

## RESEARCH DESIGN AND METHODS

### Mothers and offspring(Figure 1)

**Figure 1.**
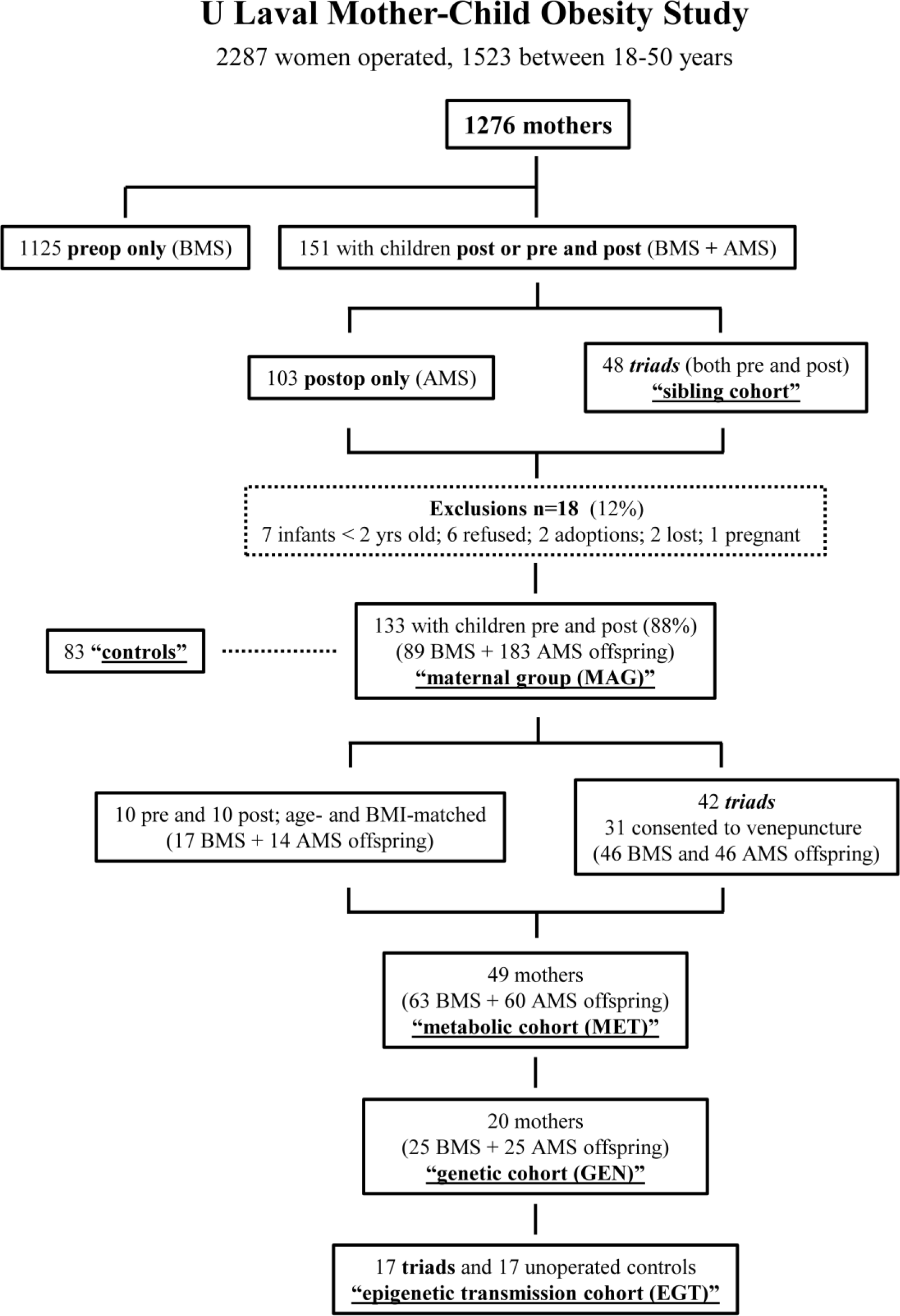
Schematic representation of the University Laval Mother-Child Obesity Study.

#### Mothers

Since 1984 we performed BPD in 2 287 consecutive, severely obese women, 1 523 aged 18-50 years in M-COS. Among these, 1 125 had given birth BMS and 103 AMS and 48 had done so both before and after surgery giving rise to triads consisting of siblings born BMS and AMS (“<u>sibling cohort</u>”). We recruited all mothers with offspring older than two years of age born AMS living within 250 km of our center. Recruitment occurred during maternal office visits, with letters and phone calls as needed. Pregnancy data on GWG, complications and birth weights were self-reported during office visits and supplemented by official hospital records (17). A cohort of 83 preoperative contemporaneous age-matched women served as “maternal <u>controls</u>” (Table 1). Of the 151 mothers giving birth AMS, 133 (88%) agreed to participate with 89 BMS and 183 AMS consenting offspring (“maternal group”; MAG).

**Table 1.**
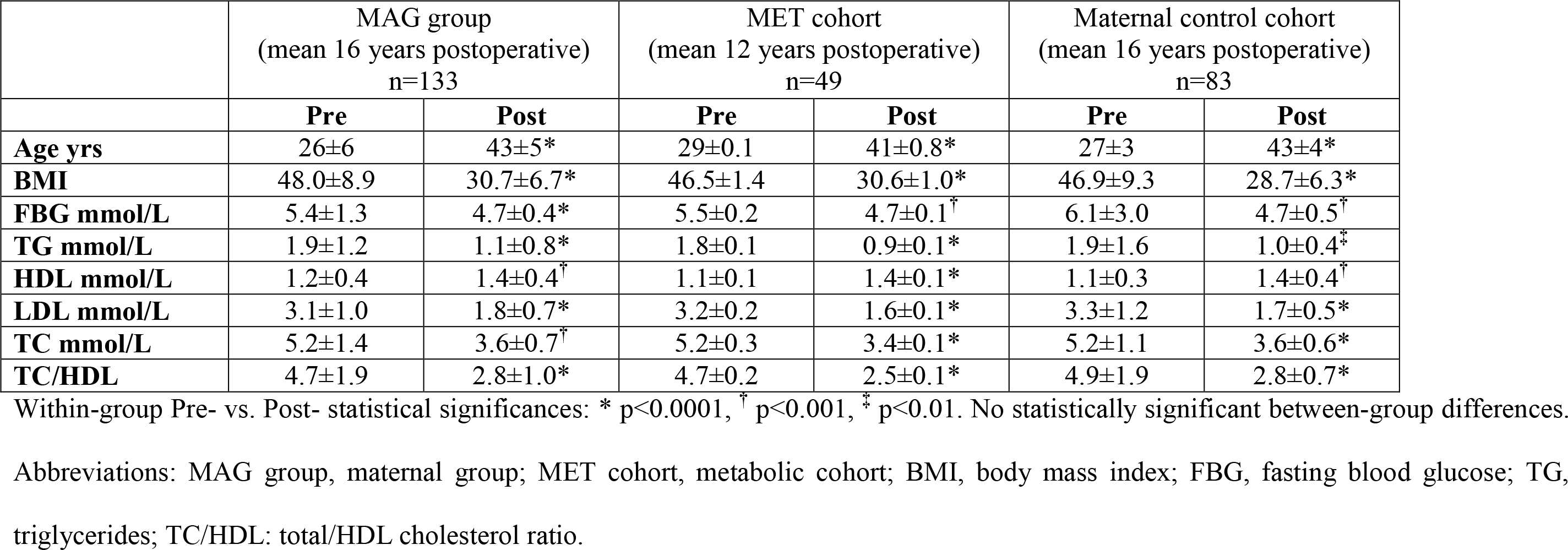
Cardiometabolic risk factors before (Pre) and after (Post) bilio-pancreatic diversion in matched operated and control women (mean ± SD).

#### Offspring

A subset of 63 BMS and 60 AMS offspring among the 89 BMS and 183 AMS underwent more extensive metabolic testing (“<u>metabolic cohort”; MET</u>) (13) and a subset of 25 BMS and 25 AMS offspring underwent genetic studies (“<u>genetic cohort”; GEN</u>) (14,15).

We selected the 17 full triads consisting of mothers with corresponding BMS and AMS offspring for further gene methylation analyses. Unoperated control severely obese women were matched to operated mothers from triads to obtain a group (“<u>epigenetic transmission cohort”; EGT</u>) (16) consisting of pre-operative maternal controls, operated mothers and their AMS and BMS offspring (17 quartets = 68 individuals).

All studies were approved by the Québec Heart and Lung Institute Ethics Committee and conformed to the Helsinki Declaration of Subjects’ Rights. Written informed consent was obtained from adults and parents and assent from minor offspring before inclusion in the study.

### Operation

Between 1984 and 1990 fifteen percent (15%) of mothers were operated with Scopinaro’s BPD, whereas after 1990 all patients had the pylorus-preserving BPD with "duodenal switch" (Supplemental Figure S1). Both procedures are based on the same mechanisms: 1) transiently limited gastric capacity, 2) sustained decreased lipid absorption owing to reduced lipase and diverted bile, 3) improved glucose metabolism from enhanced incretin release and 4) increased satiety from relatively undigested food emptying directly into the ileum bypassing duodenum and jejunum, potentiating anorexigenic peptide release. All mothers routinely took multivitamins including folate and minerals; blood levels were regularly monitored. Morbidity and mortality of BPD in our hands is comparable to that of series of other diversionary bariatric operations in the literature (6).

### Household eating habit questionnaire

Using a semi-structured questionnaire, we asked mothers 5 questions about postoperative changes in family and personal eating/shopping habits on different occasions during their postoperative office visit interviews, entered into our medical records.

### Blood chemistry

Details of physical examination, anthropometry, blood analyses etc. in subsets of offspring are published (12,13,15). FBG and plasma lipid analyses (TC, LDL, HDL, TG) were performed in the hospital clinical laboratory using standard methods. Plasma insulin, CRP, adipokines and various gastro-intestinal peptides were measured in a specialized research laboratory (13). Insulin resistance was calculated via HOMA-IR (18). Owing to the mean age difference of 8 years between the two offspring groups (BMS = 16±6 years and AMS 8±4 years; p<0.0001) all anthropometric comparisons used age- and sex-adjusted norms from national surveys (19–21).

### Anthropometry and body composition

We defined pediatric obesity using age- and sex-adjusted norms proposed by different international agencies expressed either as BMI percentile or BMI z-score: BMI percentile (<5 = “underweight”; ≥5 to <85 = “normal weight”; ≥85 to <95 = overweight”; ≥95 to <98 = “obese”; ≥98: “severely obese”; BMI z-score (<−1= underweight; ≥−1 to <1= normal weight; ≥1 to <2= overweight; ≥2 to <3= obese; ≥3= severely obese) (20,21). In the MET cohort we performed bioelectric impedance analysis (Tanita®; Arlington Heights, IL) for determination of percent body fat (19). Abdominal obesity (adiposity) was defined as WC/HT ratio exceeding the 90th percentile adjusted for age and sex in subjects with >30 percent body fat.

### Definitions

Diabetes in adults was defined as FBG ≥7 mmol/l or treatment by medication and/or diet. All pregnant women in Quebec are screened for GDM using the Canadian Diabetes Association criteria. Dyslipidemia was defined as TC/HDL>4 and plasma TG>1.7 mmol/l or on-going medication, whereas the diagnosis of hypertension was solely based on use of anti-hypertensives. We defined “metabolic fitness” based on presence of any of the established elements of the metabolic (or dysmetabolic) syndrome. Complete remission was defined as discontinuation of medication and presence of laboratory reference or “normal” levels of metrics.

### Methylation profiles

Whole genome studies of blood DNA from the EGT cohort were published previously (14–16). In families with multiple offspring born before and/or after maternal surgery the latest before and earliest after maternal surgery were chosen. Methylation levels were extracted for more than 485 000 CpG sites at a genome-wide level without regard to the presence of potential polymorphic CpG sites. Pearson correlation coefficients were calculated for methylation levels and for differences in methylation levels in mothers vs. controls and AMS vs. BMS offspring. The data reported in this paper have been deposited in the Gene Expression Omnibus (GEO) database, www.ncbi.nlm.nih.gov/geo (pending accession number).

### Statistics

Qualitative data are presented as percentages and quantitative data as mean ± SD. Differences in proportions were tested with Chi^2^ or Fisher’s exact test, and continuous variables were compared with Student’s *t* test or the Wilcoxon test. Some variables were investigated for a log-transformed eligibility to stabilize variance and corresponding p-values are for transformed values. Dependent variables were compared between and within BMS and AMS children using a repeated mixed model and adjusted for age when appropriate.

Univariate normality assumptions were verified with Shapiro-Wilk tests. Brown and Forsythe’s variation of Levene’s test statistic tested the homogeneity of variances. Data were analyzed using the statistical package SAS 9.2 (SAS Institute Gary, NC), statistical significance being defined as p≤0.05 with p≤0.15 as a trend.

The RCircos® package (22) was used to visualize differences in methylation levels (DiffScores) comparing mothers to controls, AMS to BMS offspring, and to schematize annotated CpG sites and correlation of changes in methylation levels (mothers-controls vs. AMS-BMS offspring).

## RESULTS

### Household conditions

According to our office interviews with mothers from the MAG group there were: 1) no significant differences in performance or duration of breastfeeding (BMS: 40%; AMS: 34%) 2) no offspring on diets in either group, 3) no changes in size or content of family meals after surgery among 75% of mothers; 58% reported not having changed the content of their own diet. 4) Socio-economic conditions were relatively unchanged and 5) BMS and AMS children lived in the same family household for approximately 60% of their lives.

### Mothers and Offspring

Baseline prevalences of diabetes, dyslipidemia and hypertension of the 133 MAG group mothers were similar to those of the 83 matched pre-operative “control” women by design (Table 1). Postoperatively, 12 years after the operation, mothers from the MET cohort had 16 units lower BMI. During their first postoperative pregnancy, 4.0±1.7 years after their surgery, GWG was 67% lower than before surgery (4.6±9.4 vs. 13.9±13.9 kg before; p<0.0003). Postoperative mean fasting blood glucose (FBG) was 15% lower (p<0.001). Plasma insulin and homeostasis model assessment (HOMA-IR), available solely for postoperative mothers, were normalized (mean insulin 14.8±15.5 μU/mL; HOMA-IR 2.54±1.38) (23,24) and measures of preoperative dyslipidemia, [total to high density lipoprotein cholesterol ratio (TC/HDL)>4 and plasma triglycerides (TG)>1.7 mmol/l] decreased after surgery by 88 and 80 % respectively (p≤0.0001) (Table 1). The 15% of women with pre-operative hyperglycemia (>6 mmol/l) and 20% with diabetes were in complete remission. Clinical results in the 20 mothers in the epigenetic cohort were similar (16).

There were large differences in cardiometabolic risk factors between BMS and AMS offspring from the MET cohort (Table 2) exhibiting sexual dimorphism of improvements in AMS offspring. In BMS boys prevalences of severe obesity and dyslipidemia were statistically significantly greater than girls. In the total offspring cohort, BMS adolescents (14 males, 19 females, aged 17±2 years) exhibited more obesity, severe obesity (BMI z-score; p<0.05) and adiposity [waist circumference to height (WC/HT) ratio; p=0.03] than AMS offspring (7 boys, 9 girls, aged 16±2 years). In aggregate AMS adolescents displayed improved insulin sensitivity, dyslipidemia, and lower C-reactive protein (CRP) than BMS. In contrast, cardiometabolic differences increased with age comparing pre-pubertal children to adolescents. Thus, these differences appeared to take time to be expressed after puberty: most elements of the metabolic syndrome were improved compared to before age 10 (Supplemental Table S1).

**Table 2.**
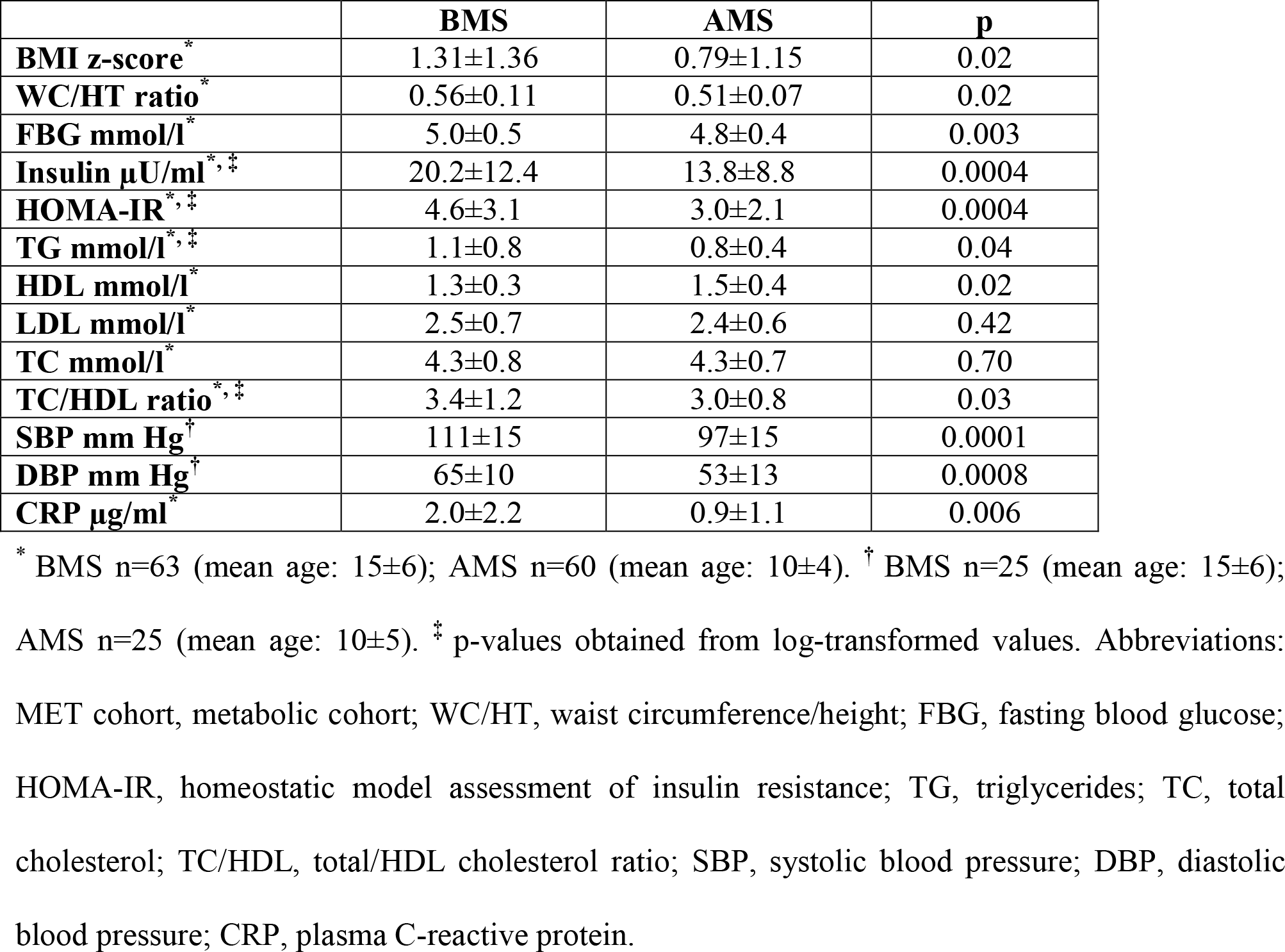
Cardiometabolic risk factors in offspring born before (BMS) and after (AMS) maternal surgery (MET cohort; mean ± SD).

### Intergenerational correlations

The difference in prevalence of severe obesity between preoperative control women and postoperative mothers was 77% versus 53% between BMS and AMS offspring, associated with lower plasma insulin levels. The insulin levels were 81% lower in mothers than controls and 30% lower in AMS than BMS offspring consistent with lower HOMA-IR: 82% lower in mothers than controls and 31% lower in AMS versus BMS offspring.

In the MET cohort, mother/offspring correlations BMS were significant for glucose (r=0.41, p=0.0009, insulin not measured) and AMS for glucose (r=0.33, p=0.01, insulin r=0.44, p=0.002; and HOMA-IR r=0.37, p=0.01). Further details on correlations according to maternal status are provided in Supplemental Table S2.

### Trajectories of dysmetabolic markers

Weight increased more rapidly with age in BMS than AMS offspring from the MAG group, exhibiting a correlation of BMI z-score vs. age of r=0.33 in BMS (n=89 observations; p=0.002) while almost negligible in AMS: r=0.10, (n=183 observations; p=0.20) corresponding to a 3-fold faster weight rise in BMS offspring [slope: 0.099 for BMS vs. 0.036 for AMS offspring (p=0.13)], a statistical trend. The difference was significant for girls (p=0.02) but not for boys. In offspring aged 2-26 years from the MET cohort, abdominal obesity (WC/HT) also increased significantly faster in BMS (n=62; slope 0.008) than AMS (n=60) in whom the slope was negligible (0.001) with group between-slope difference: p=0.04 (Figure 2), discordant for girls (p=0.054) vs. boys (p=0.40). Between-slope differences for TG (p=0.11), TC (p=0.14) and TC/HDL ratio (p=0.07) exhibited trends also varying between boys and girls. Correlations between BMI z-score and cardiometabolic markers consistently demonstrated higher correlations in BMS than AMS, entailing lower risk associated with obesity in AMS offspring (Supplemental Figure S2).

**Figure 2.**
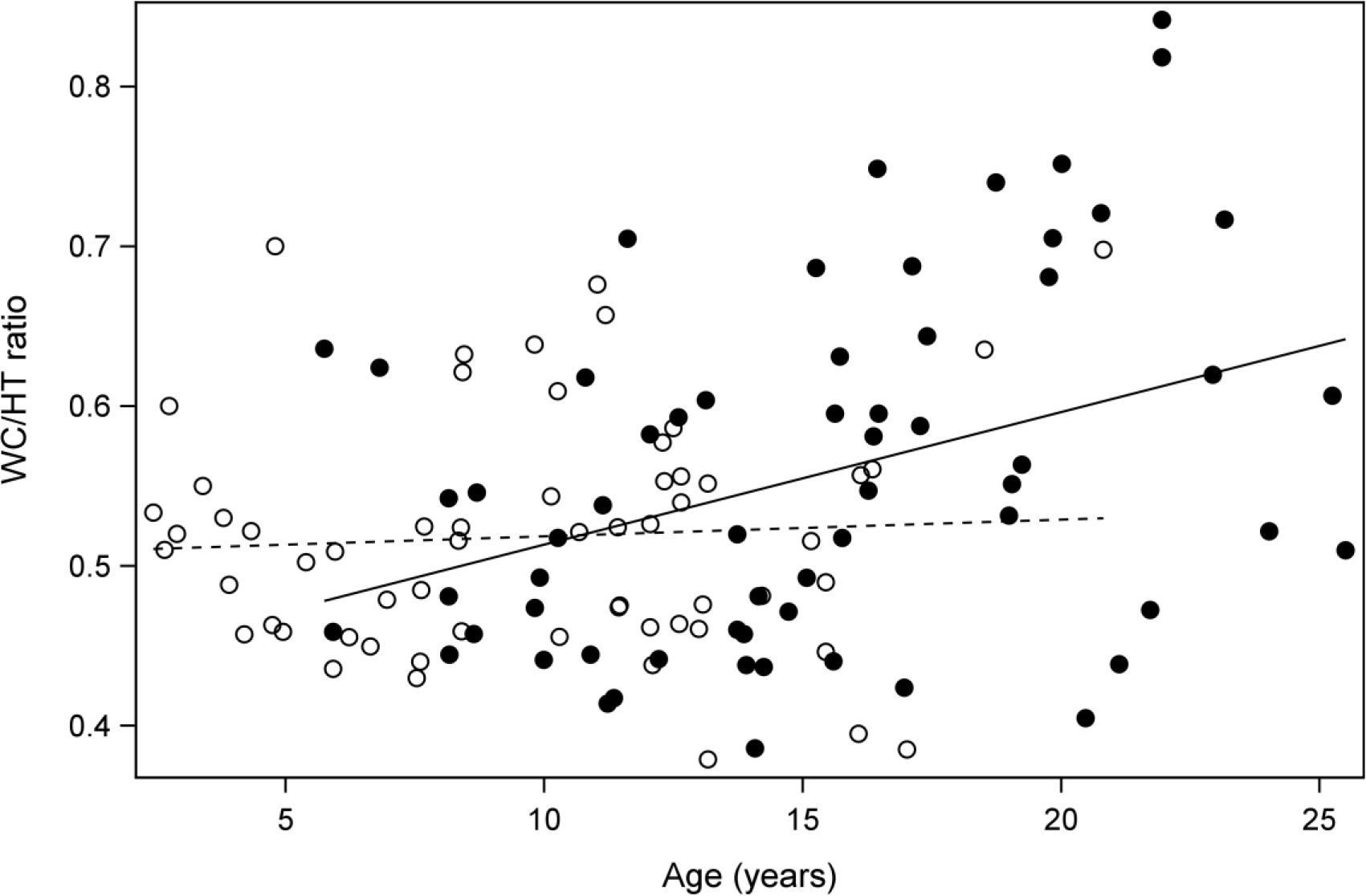
Age trajectories of abdominal obesity. Age trajectories of abdominal obesity (WC/HT; waist circumference/height) in BMS (filled circles; n=62) slope (solid line; 0.008±0.003) *vs.* AMS (empty circles; n=60) slope (dashed line; 0.001±0.002) in offspring aged 2-26 years (between-slope difference: p=0.04) from the MET cohort.

### Methylation profiles in offspring

Our findings of intergenerational correlations (Table 3) prompted us to search our methylation data (14–16) for differentially methylated CpG sites in AMS vs. BMS offspring concordant with differences between mothers and controls. To integrate methylation data, we used a liberal cut off (|DiffScore|≥13 = p≤0.05) in our EGT cohort (“quartets”). Mothers and controls had 321 347 differentially methylated sites whereas siblings (AMS vs. BMS) only had 45 625. Most of the differentially methylated sites in mothers and controls had lower methylation levels in the mothers reflected in a higher prevalence of negative DiffScores (Supplemental Figure S3). Counter-intuitively, the majority of differentially methylated sites between AMS and BMS offspring demonstrated higher methylation levels in AMS offspring although the differences were in the lower range. DiffScores varied from −371 to 374 in mothers vs. controls (mean −46.2±47.8) and from −46 to 56 in AMS vs. BMS (mean 12.5±12.5). We identified 33 232 common differentially methylated sites. Correlation analyses for CpG sites showing directionality in methylation differences (4 446 sites) revealed 1846 sites with positive mother-control and AMS-BMS correlations among which 154 were moderate or strong (r≥0.4). Pearson’s correlation coefficients for these 154 CpG sites are depicted in Figure 3 (outer circle) along with gene names for the 122 sites assigned to 120 unique genes. Gene enrichment analysis from the list of correlated CpG sites revealed 15 overrepresented pathways among which: 1) cell growth, proliferation and development, 2) organism growth and development, 3) amino acid metabolism, and 4) lipid and carbohydrate metabolism (Supplemental Figure S4).

Correlations between methylation levels and metabolic analytes were assessed for CpG sites assigned to genes from overrepresented pathways related to amino acid, lipid or carbohydrate metabolism as well as for those from genes associated with cardiometabolic risk profile in genome-wide association studies (full list provided in Supplemental Table S3). Notably, methylation levels of CpG sites located within *INS-IGF2* loci and TG levels and *A1BG* methylation and FBG levels were significantly correlated. Methylation levels of the protein kinase AMP-activated non-catalytic subunit gamma 2 (*PRKAG2*) gene, assigned to the PXR/RXR activation pathway, were negatively correlated with plasma lipids (TC, LDL-C, TG, TC/HDL) and insulin levels. Furthermore, correlations between methylation levels and BMI were identified in mothers and controls for 7 of these sites (Supplemental Table S4) while methylation levels of a site located in carbohydrate sulfotransferase 11 (*CHST11*) was correlated with BMI percentile in offspring (r=0.368, p=0.038).

## CONCLUSIONS

We have previously shown that improved maternal metabolic fitness after bilio-pancreatic bypass, a lipid-sequestering metabolic ‘bariatric’ operation, is associated with improved pregnancy outcomes (17) and fewer cardiometabolic risk factors in AMS offspring compared to BMS siblings (13) associated with differential methylation of inflammatory and glucoregulatory genes suggesting the presence of epigenetic mechanisms malleable in the intrauterine environment (15). Subsequently, differential methylation of inflammatory and type 2 diabetes genes has also been reported in AMS and BMS siblings (25) supporting the implication of such mechanisms. Here we extend these observations by uniquely demonstrating differential trajectories for cardiometabolic factors favoring AMS offspring vs. BMS siblings, and by detecting maternal-offspring correlations commensurate with differential methylation between mothers and matched un-operated control women.

The implications are that improving maternal fitness before and/or during gestation has durable beneficial effects on the offspring with the potential to abrogate generational transmission of the dysmetabolic diathesis in children of obese parents (1). Specifically, offspring born after maternal bariatric demonstrated improved cardiometabolic profiles in comparison to BMS siblings here and in various cohorts (12,13,26). There is a remarkable resemblance between the steep trajectories of BMI z-scores and central obesity among our BMS offspring aged 2-26 years and the simulated (to 35 years) and observed (10 year data) trajectories of childhood obesity into adulthood published in 2017 (10). Those analyses derived from large US databases failed to consider sex differences. Our results suggest a sexual dimorphism in trajectories of BMI z-score with age, differences between AMS and BMS being identified in girls only. Abdominal obesity (WC/HT) was also shown to increase faster in BMS than AMS (Figure 2), an effect mainly driven by a difference observed in girls. Such observations may be partially explained by the limited number of adolescent boys in our cohort (21 adolescent boys vs. 28 adolescent girls) or by the relatively young age of our cohort. Larger differences in cardiometabolic markers between AMS and BMS adolescents than pre-pubertal children suggest that differences in trajectories and cardiometabolic markers may take time to express. Such observations from our group and others (26), combined with the age of our cohort, may partially explain sexual dimorphism observed here without obviating the potential contribution of puberty (27), sex hormones and other biological factors (28) in the diathesis of cardiometabolic risk profile. Nevertheless, our findings suggest that metabolic operations in reproductive-age women, by normalizing BMI z-score trajectories in offspring born after maternal BPD, have the potential to curb the burgeoning epidemic of adult obesity (10,29) and type 2 diabetes (11).

The concordance between differential methylation of cardiometabolic genes in offspring and operated mothers suggests specific epigenetic mechanisms, similar to those recently found by others in sperm before and 1 year after metabolic gastric bypass surgery (30), as well as in muscle tissue 6 months after gastric bypass (31) where we now uniquely demonstrate the durability of epigenetic markers extending over 26 years in offspring. Interestingly, a CpG site located within the promoter region of the poly(A) binding protein cytoplasmic 4 (*PABPC4*) gene, herein correlated with cardiometabolic risk factors and BMI, was previously found to be differentially methylated between obese and lean men (30). Our approach proposes potential targets for the transmission of improved maternal fitness to offspring revealed in correlations of methylation levels. For instance, we identified an inverse correlation between methylation, lipid and insulin levels for a CpG site mapped at the *PRKAG2* loci. This gene encodes a regulatory subunit of the AMP-activated protein kinase, an energy-sensing enzyme regulating *de novo* biosynthesis of fatty acid and cholesterol, and was previously shown to be associated with diabetes incidence (32) thus providing a potential target for further studies of cardiometabolic risk transmission to offspring.

There are several putative explanations for our findings related to proposed mechanisms for maternal transmission of the dysmetabolic diathesis. Hyperglycemia, insulin resistance and dyslipidemia are related to inflammatory teratogenic changes affecting the fetus. Placental function is also impaired in diabesity related to both gluco- and lipotoxicity, with adverse effects on the fetus. The uniquely lipid-sequestering BPD operation lowers free fatty acid levels and abrogates the lipotoxicity engendering insulin resistance, islet cell failure and mitochondrial dysfunction affecting efficiency and immune function of the placenta (33) as well as fetal tissues. Beta-cell failure of maternal diabesity in the form of GDM or ‘pre-diabetes’ transmitted to BMS offspring is likely reversible in mothers over time (34), potentially explaining our finding of improved glucoregulation in AMS offspring compared to their BMS siblings.

Collateral benefits of maternal BPD, beyond preventing GDM, hypertension and eclampsia, primary or recurrent (13,17) are: reducing the prevalences of macrosomic, large for gestational age births and caesarian delivery (35,36). Small for gestational age (SGA) births were increased postoperatively in our population, as is the case in general after ‘bariatric’ surgery (7), but there was no increase in premature births in either study. Although severe obesity increases risk of premature deliveries, implicit in C-section for a macrosomic fetus, and there were no statistically significant increases in premature deliveries after bypass operations, care must be taken to avoid over-interpretation of these data recognizing that long term neurological risks of SGA and premature deliveries require more research.

There are several limitations in our study. Life-style and household findings are based on retrospective self-reports, although consistent with postoperative recommendations and studies demonstrating high caloric intake after BPD (37). Indeed, the BMS offspring sharing the same household did not exhibit any changes in *habitus* after maternal surgery (13). The questionnaires and interviews were not sufficiently detailed to evaluate potential dietary differences.

The size of this patient series might appear small. Because of the rarity of bariatric surgery, especially BPD before the last 10-15 years, our patient series tracked over 25 years is exceptionally large with a high follow-up rate. Although severely obese mothers electing to undergo bariatric surgery represent a select subgroup among severely obese patients, any such differences were present before maternal surgery and during earlier pregnancies and our control women were drawn from preoperative BPD patients. We lack data on paternal fitness, recently recognized as transmissible via sperm and influenced by paternal surgery (30).

In the absence of precise Tanner stages for the children, we arbitrarily chose age 14 as cut-off for “post-pubertal”. A recent metabolomic study of associations between mothers and their 6-10 year-old children did not detect any differences in relevant metabolites attributable to presence or absence of pubertal signs present in 30% of the children (36).

Our study was not designed or powered to follow sufficient numbers of offspring to present longitudinal data. Such studies would be prohibitively expensive requiring large multi-center participation to recruit adequate numbers of randomly occurring pregnancies and births with subsequent prolonged follow-up. The 26-year time period is long enough for relevant developmental observations yet short enough to preclude major changes in secular trends in Nutrition and Maternal-Fetal-Medicine.

Although the practice of bariatric surgery has relatively recently incorporated less invasive laparoscopic approaches, our operation is part of current practice and virtually unchanged over this period. We did not detect in either cohort or sex any discontinuity in the dysmetabolic parameters which would indicate differences in rearing environment or childhood experiences (38). Although our observations are valid in this cross-sectional design they might not be applicable to mechanistically different operations with inferior or different maternal outcomes.

Our study cannot determine the timing of the epigenetic changes although others have shown them to emerge relatively early postoperative (39). Nonetheless, a slightly larger study on offspring from mothers who underwent various different types of bariatric surgery identified similar magnitude of differences in methylation levels (∼23 500 CpG sites) between AMS and BMS siblings aged 6-33 years (25). Here we show that epigenetic changes are durable, exhibiting ‘metabolic memory’, both in mothers studied mean 16 years postoperative and in AMS offspring assayed in adolescence. This is an important finding contrasting with the traditional genetic determinism: our surgical study joins others employing life-style measures and drug treatment in demonstrating the plasticity of epigenetic changes over the lifecycle. Accordingly, methylation-based age estimation (“biological age”) has now been associated with cardiovascular diseases (40).

Although un-realistic on a population level owing to scarcity of requisite resources, by improving parental fitness durably transmitted to subsequent offspring, metabolic surgery has the potential to significantly abrogate continuing epidemic spread of the dysmetabolic syndrome by creating a new, beneficial “metabolic memory”. Obviously our small study does not allow population-level conclusions. It provides mechanistic insights generating testable hypotheses regarding the impact of modifiable precursors of the inflammatory dysmetabolic syndrome of adiposity.

We conclude that beneficial metabolic effects after BPD in severely obese mothers improved the intrauterine environment of subsequent pregnancies exhibiting normalized GWG and metabolically fit offspring. Ensuing changes at the DNA methylation level many years after birth were associated with decreased cardiometabolic risk, potentially improving quality adjusted life years compared to offspring born before maternal BPD surgery. Maternal metabolic surgery has pre-emptive preventive potential to abrogate development of obesity and type 2 diabetes in future generations.

## Author Contributions

PM initiated the U Laval Mother-Child Obesity Study and operated and managed all patients together with SM and the surgical team, accruing all clinical data. PM and JGK designed the study making intellectual contributions, conceiving the analyses and interpretations of data, ultimately writing the paper together with FG and M-CV who performed the critical laboratory assays. SS and FG did statistical analyses. All authors approved the final version. M-CV is the guarantor of this work and, as such, had full access to all the data in the study and takes responsibility for the integrity of the data and the accuracy of the data analysis.

## Acknowledgments

Our studies could not have been performed without major contributions of the bariatric surgeons: Simon Biron MD, Frédéric-Simon Hould MD, Stéfane Lebel MD, Odette Lescelleur MD and Laurent Biertho MD and our dedicated Research Assistant: Paule Marceau BA. We appreciate critical review with constructive editorial advice from Professor Bruce S. McEwen, PhD, The Rockefeller University. This study was supported by MCV Canadian Institutes of Health Research (CIHR) and Heart and Stroke Foundation of Canada (HSFC) grants as well as Department of Surgery funds. MCV is Canada Research Chair in Genomics Applied to Nutrition and Health.

## SUPPLEMENTAL FIGURE LEGENDS

**Supplemental Figure S1. Schematic representation of biliopancreatic diversion with duodenal switch.** Biliopancreatic diversion operations (BPD) performed in all women: 15% BPD (Scopinaro’s distal gastrectomy) [21] and 85% BPD-DS (sleeve gastrectomy with pylorus-preserving “duodenal switch”).

**Supplemental Figure S2. Relationships between obesity (BMI z-score) and cardiometabolic risk factors in BMS (filled circles/squares, n=63) *vs*. AMS (empty circles/squares, n=60) from the MET cohort.** Slopes of correlations in BMS (solid lines) vs. AMS (dashed lines) offspring for TG (Panel A; slopes 0.302±0.06 vs. 0.136±0.047; p=0.04), TC/HDL ratio (Panel B, slopes 0.534±0.096 vs. 0.289±0.087; p=0.07) and WC/HT ratio (Panel C, slopes 0.070±0.006 vs. 0.051±0.005; p=0.016). Panel D shows slopes of WC/HT ratio in BMS and AMS offspring according to sex: BMS males (filled squares, solid line; slope 0.070±0.008) vs. AMS males (empty squares, short dashed line; slope 0.054±0.009; p=0.24) and BMS females (filled circles, long dashed line; slope 0.071±0.008) vs. AMS females (empty circles, dashed line; slope 0.050±0.006; p=0.055).

**Supplemental Figure S3. Circos plot of methylation differences in mothers vs. controls and AMS vs. BMS offspring and correlated CpG sites.** Circos plot showing (from inner circle to outside circle) chromosome ideogram with chromosome number (1), heat maps of DiffScores representing differences in methylation level between mothers and controls (2) as well as between AMS and BMS offspring (3) for commonly differentially methylated CpG sites (n=33 232). Pearson’s correlation coefficients of the 154 CpG sites show changes in methylation levels (r>0.4; mothers-controls vs. AMS-BMS offspring) (4) along with corresponding gene annotations (5).

**Supplemental Figure S4. Pathways enriched from the list of 154 CpG sites with correlations between differences in methylation levels in mothers-controls and AMS-BMS offspring.** Pathways related to cell (red) and organism growth and development (green) and amino acid (blue), lipid (purple) and carbohydrate (orange) metabolism. Number and percentage of genes identified for each pathway are shown above bars.

